# A bioinformatics pipeline for the assessment of the evolutionary relationship of some *Drosophila* species based on class II transposons mapping

**DOI:** 10.1101/2022.09.13.507812

**Authors:** Nicoleta-Denisa Constantin, Alexandru Marian Bologa, Attila Cristian Ratiu, Alexandru Al. Ecovoiu

**Affiliations:** University of Bucharest, Faculty of Biology, Department of Genetics, Intrarea Portocalelor Street, No. 1-3, 060101, Bucharest, Romania

## Abstract

Transposons are mobile DNA sequences, known for their ability to insert into other locations in the genome. Genome sequencing allowed the identification of the high content of transposons in various model organisms. Here, we present a bioinformatics pipeline developed to estimate the evolutionary relationship between *Drosophila melanogaster* and other *Drosophilidae* based on the comparative analysis of the presence and distribution of class II transposons. Our study reveals that the presence and distribution of transposons hobo, HB, Tc1, Tc1-2, hopper and Bari1 points to close evolutionary relationship among *D. melanogaster, D. simulans, D. sechellia* and *D. yakuba*, which is in accordance with other data available in literature.

## Introduction

Consecutive to the sequencing of the genomes of different species, the transposons were identified either as integral sequences or as incomplete sequences (transposon fragments), which we will hereafter call residual sequences or remnants. Often, nucleotide sequences of transposon fragments are identical, but their location in the genome can be preserved or variable.

In bioinformatics, there are two different practical approaches, the use of browser-based tools that are easily accessible, such as databases or applications. On the other hand, there is the command-line tools option that offers maximum flexibility, precision, speed and efficiency for large data set analysis (Pevsner, 2015). We used both procedures to obtain a pipeline that performs a comparative analysis of class II transposons in the genomes of 12 species of *Drosophilidae*, in order to display evolutionary relationships among these species.

In this study, we sought to map the remnants of the following class II transposons: hobo, HB, Tc1, Tc1-2, Bari1, Bari2, Helitron, looper1, hopper and hopper2. The standard/model species is *Drosophila melanogaster*, and the species of interest are *D. sechellia, D. simulans, D. yakuba, D. erecta, D. ananassae, D. pseudoobscura, D. persimilis, D. willistoni, D. mojavensis, D. virilis* and *D. grimshawi*.

## Implementation

Our bioinformatics pipeline consists of four original scripts, plus a few open-access existing programs and contains the following five modules: processing the *Drosophilidae* genomes (previously downloaded); processing the residual transposon sequences (downloaded from *FlyBase*); extraction of genome-transposon junction sequences; quality check for the alignments of the nucleotide sequences (with Kablammo) and the comparative analysis of the preservation of genome-transposon junction sequences with Genome ARTIST (Figure 1).

**Figure 1.**
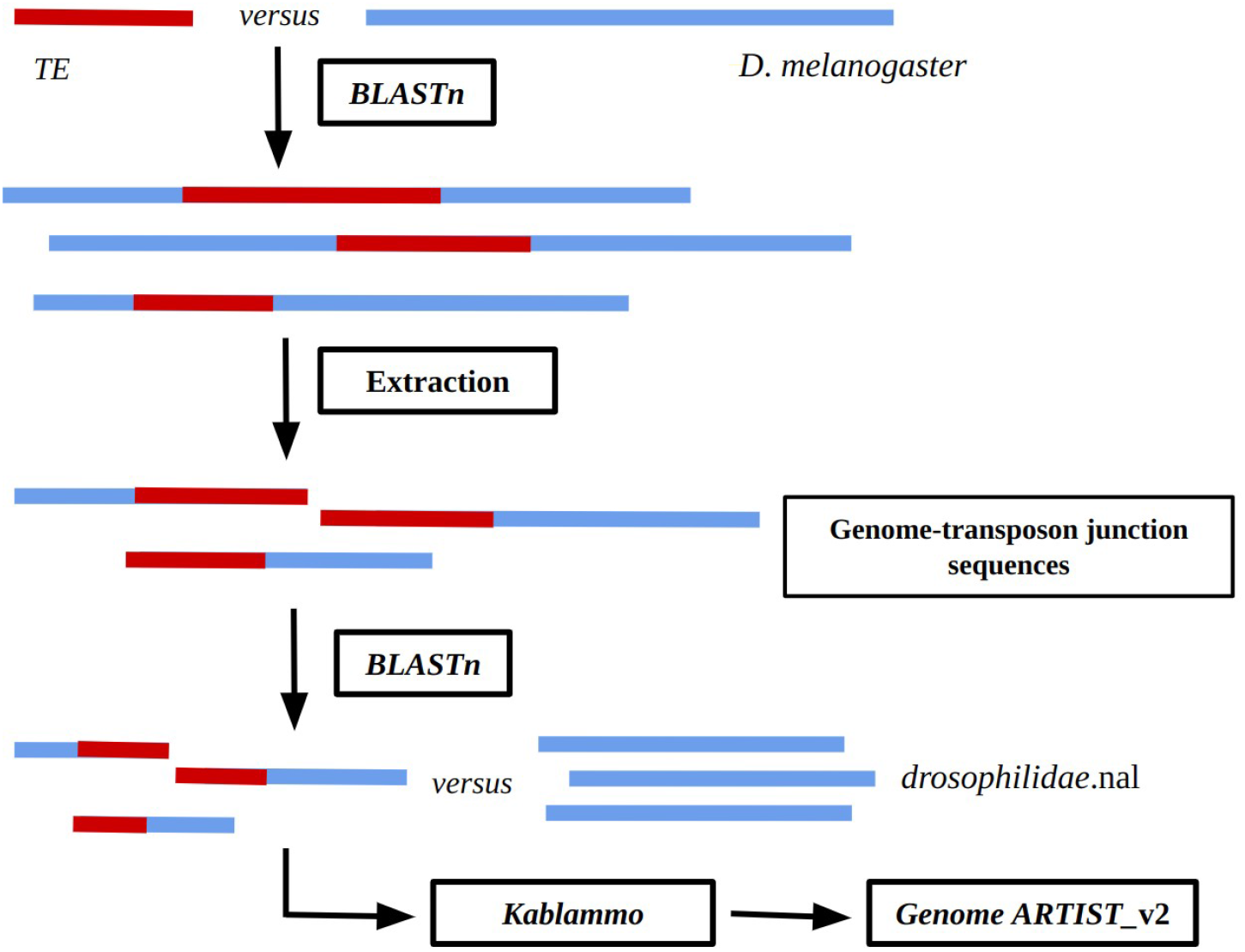
The main modules of the bioinformatics pipeline that allows for transposon identification in available *Drosophilidae* genomes. Genomic sequences are represented by using blue horizontal lines, while transposon specific sequences are depicted with red lines.

### 1. Downloading and processing the *Drosophilidae* genomes

The genomes of interest have been downloaded from the *FlyBase* database as multi-FASTA files. The genome processing script (*1_genome.sh*) modifies each header by appending the name of the species; in addition, it creates specific *BLASTn* databases for each of the 12 genomes, and defines an alias, a local database called *drosophilidae*, containing 11 genomes, excepting the *D*. *melanogaster* standard genome.

### 2. Downloading and processing the residual transposon sequences

The residual transposon sequences were downloaded from *FlyBase* in a *multi-FASTA* format and processed according to the substepts shown in Figure 2. The *2_remnants_processing.sh* script involves dividing the *multi-FASTA* file into several single *.fa* files (one for each residual transposon sequence existing in *FlyBase* file). The user specify the acronym of a transposon (i.e for hobo transposon, one should type its acronym, “H{}”) and the files that do not contain the acronym of the fragment of interest in the name are deleted. The remaining. *fa* files are titled according to the name of the residual sequence and the chromosome in which it is identified (for example, for the remnant H{}84 sequence of the hobo transposon, the file name is “H{}84-X.fa”).

**Figure 2.**
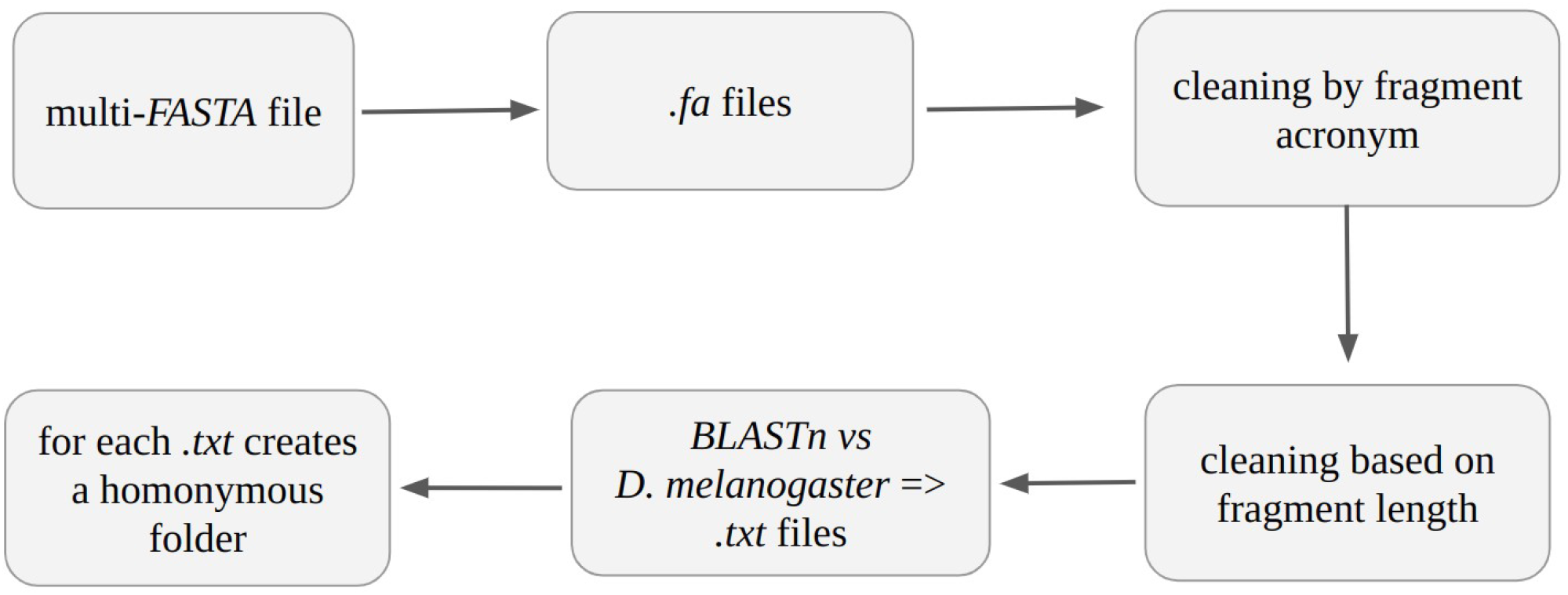
The main steps of the processing of residual sequences.

In this study, residual sequences of at least 30 nucleotides were analyzed, which is why we implemented a filtration step, which eliminates shorter sequences than the specified threshold.

The next step is to align the residual sequences versus the *D*. *melanogaster* reference genome, with *BLASTn;* the results are stored in a tabular format, in *.txt* files of the same name. This stage is necessary to identify the location of the transposon fragments in *D*. *melanogaster* genome (genomic coordinates).

The last step is to organize the resulting files. For each *txt* file, a folder with the same name is created and it contains the corresponding file. Thus, at the end of this step, there will be available a collection of the fragments to be analyzed, as well as the results of *BLASTn* alignment in separate folders.

### 3. Extraction of genome-transposon junction sequences

Genome-transposon junction sequences are extracted using *SeqKit* (Shen et al., 2016)., based on sequence coordinates range and the substeps shown in Figure 3. This interval is calculated using *3_upstream_extraction.sh* or *4_downstream_extraction.sh* scripts. When one script is running, the user is asked to specify the lengths of the sequences that will be extracted from the genomic region (genomic flanking length) and from the transposon internal region (transposon length). Based on these choices, the extraction interval is calculated.

**Figure 3.**
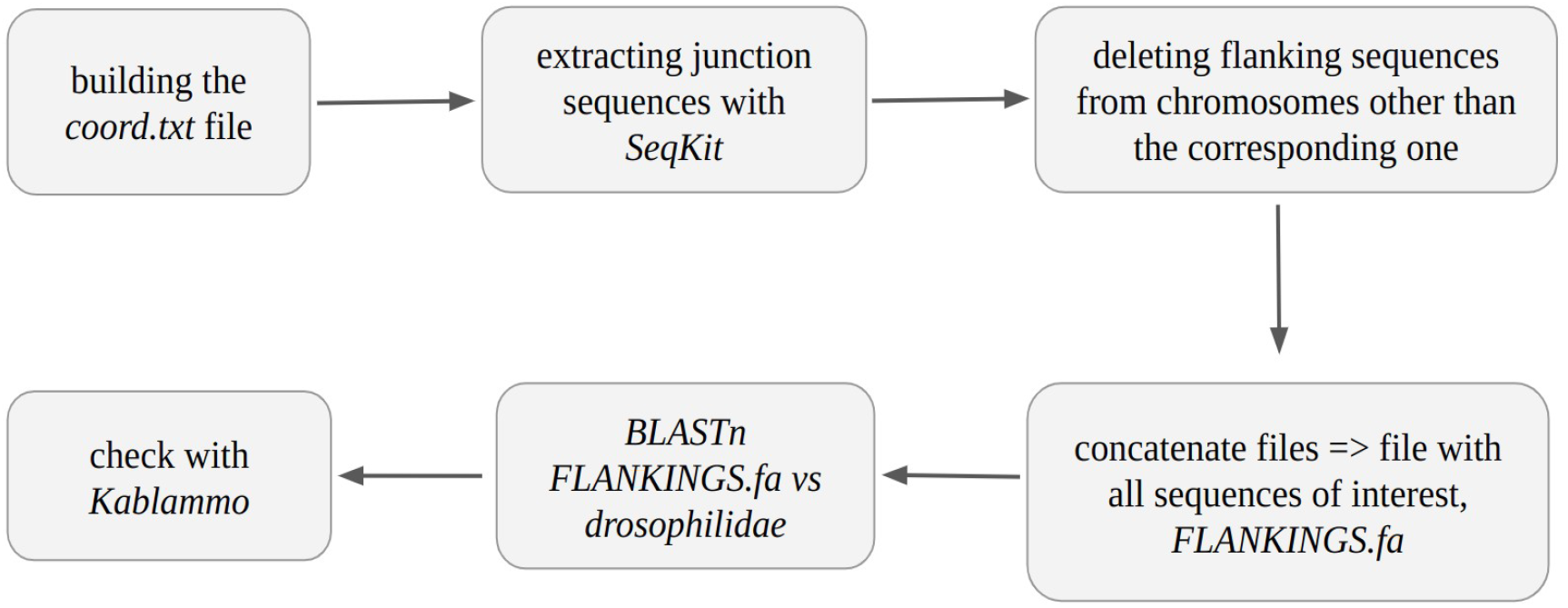
The main steps of the extraction of genome-transposon junction sequences.

Extracting the custom sequences of interest is not possible without the genomic coordinates of the transposon, so the first step is to build a file called *coord.txt*. This file contains the genomic coordinates at which the residual sequence is identified, as well as its orientation. In the header of a file containing the corresponding residual sequence is mentioned its localization in *D*. *melanogaster* genome. To build the *coord.txt* file we used the chromosome and the first coordinate from the header. These were compared with the *BLASTn* results previously obtained for the same residual sequence. A comparison of these is necessary because the version of the *D*. *melanogaster* genome (r6.43) is different from the version used by *FlyBase* for mapping residual sequences (r6.39). If these coordinates are identical, the *coord.txt* file is built.

The extraction of the sequences of interest can be performed both upstream and downstream of the genomic coordinates of the fragment, as shown in Figure 4. Therefore, by sequences of interest, we define the genome-transposon junction sequences, which contain a “blue subsequence” and a “red subsequence”.

**Figure 4.**
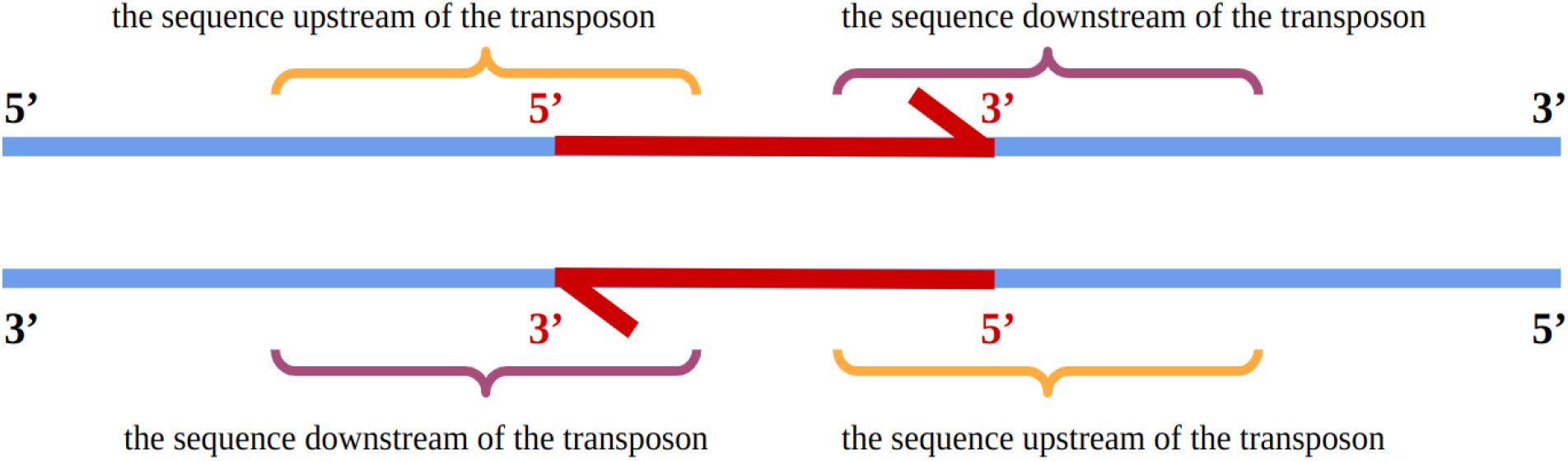
The extraction model of genome-transposon junction sequences. Color code: Red - transposon; blue - genomic DNA sequence.

In this study, we used genome-transposon junction sequences of different lengths. The length of the genomic flanking ranged from 250 nucleotides to 500 and, respectively 1000 nucleotides (nt). The transposon internal region has 30-500 nucleotides, because residual sequences that may have at least 30 nucleotides are taken into account; in these cases, we used the integral residual sequence.

The *coord.txt* file contains the location of the transposons in the reference strand. The transposons can be inserted in either the reference or in the non-reference strand of the genome (known as “-” strand). The genomic files contain only the sequences of the corresponding reference strand. Thus, for the transposon fragments that have the reference strand in non-reference strand of the genome, we have converted the junction sequences into the reverse complement strand.

*SeqKit* does not take into account the chromosome, so it extracts a subsequence defined by a particular set of coordinates from each chromosome of the *D*. *melanogaster* genome. To solve this inconvenient, we have implemented a procedure that, based on the information in the header, it removes extracted sequences belonging to chromosomes other than the one of interest.

Before concatenating the junction sequences extracted for each fragment, we append to the header of each fragment its name and the chromosome in which it is identified. The next step is to concatenate the junction sequences into a single *multi-fast*a file. Finally, several *BLASTn* alignments are perfomed between the file containing all junction sequences and the alias local database, *drosophilidae*. The results are obtained in *XML* format, so that they can be uploaded to *Kablammo*.

### 4. The Kablammo quality check for the alignments of the nucleotide sequences

Given the large number of residual sequences considered in this study, we implemented a quality check of alignments made with *BLASTn*. For this pourpose, we used *Kablammo* application, making a list of junction sequences that are to be mapped with *Genome ARTIST*.

When a particular junction sequence is used as a query, the resulting alignment must contain the junction nucleotide, at least 100 nucleotides from the transposon and at least 100 nucleotides from the genomic sequence in order to consider it for the conservation comparative analysis step. Junction sequences that provide two alignments, one of more than 100 nucleotides from the transposon and another of more than 100 nucleotides from the genome, are also accepted for the next step. In this case, the alignments must be located close to each other within a specific genome. If the sequence of the fragment of interest is less than 100 nucleotides, the junction sequence must contain it completely.

### 5. Comparative analysis of the preservation of genome-transposon junction sequences with *Genome ARTIST version 2*

The junction sequences obtained and filtered in the previous stages are loaded into *Genome ARTIST* version 2.0 (www.genomeartist.ro) for the mapping step. For further verifications of the extracted junction sequences, we use a package containing all 12 *Drosophilidae* genomes, each containing gene annotations, while the *D*. *melanogaster* genome also contains natural transposons’ annotations.

For a junction sequence extracted from *D*. *melanogaster* and identified in at least one another *drosophilid* species, we considered alignments that contain an arbitrarily threshold of 30-100 nucleotides from the residual sequence of the transposon, plus 100 nucleotides from the adjacent genomic sequence upstream or downstream of the remnant.

## Results and discussion

In this study, we analyzed the presence and distribution of ten class II transposons, respectively 230 residual sequences. For each residual sequence, we extracted six junction sequences, according to the extraction procedure, previously presented. Accordingly, we analyzed a total number of 1380 genome-transposon junction sequences.

Most preserved residual sequences were identified in the genome of *D*. *sechellia*, a smaller number in the genome of *D*. *simulans*, and the fewest in the genome of *D*. *yakuba*. For *D*. *erecta, D. ananassae, D. pseudoobscura, D. persimilis, D. willistoni, D. mojavensis, D. virilis* and *D*. *grimshawi* species we did not detect any residual sequences with preserved localization.

Between *D*. *melanogaster* and *D*. *sechellia* we identified 21 residual sequences that share the relative genomic location. Seven of these belong to hobo, and seven to HB. Bari1 has three remnants whose location are preserved between these two species, while hopper has two, and Tc1 and Tc1-2 each have one sequence.

*D*. *melanogaster* and *D*. *simulans* have in common 16 genomic local landscapes harboring residual sequences. The HB transposon is found in eight location shared by these species, while the hobo is found in only six common locations. In addition, the two species have one Tc1 and one hopper, residual sequences present in similar genomic regions.

On the other hand, when comparing *D*. *melanogaster* and *D*. *yakuba*, we identified only three conserved residual sequences, one that belongs to hopper2 and two to Tc1-2.

The results are consistent with phylogenetic data that group *D*. *melanogaster*, *D*. *simulans*, *D*. *sechellia* and *D*. *yakuba* species into the *Sophophora* subgenus, the melanogaster group, the melanogaster subgroup (Miller et al., 2021; Drosophila 12 Genomes Consortium, 2007).

By extracting and analyzing junction sequences, we identified six genes (*Tusp*, *Usp2, lncRNA:CR44997, hrm, plexA, Ada2b*) from *D*. *melanogaster* that are structural orthologous to unannotated genes from *D*. *sechellia* and *D*. *simulans*. The most interesting case (Figure 5) is that of the *Usp2* gene, which contains a residual HB in all three species. *Usp2* encodes for a hydrolase that deubiquitinates the target polyubiquitinated proteins, a process that is essential for calcium homeostasis.

**Figure 5.**
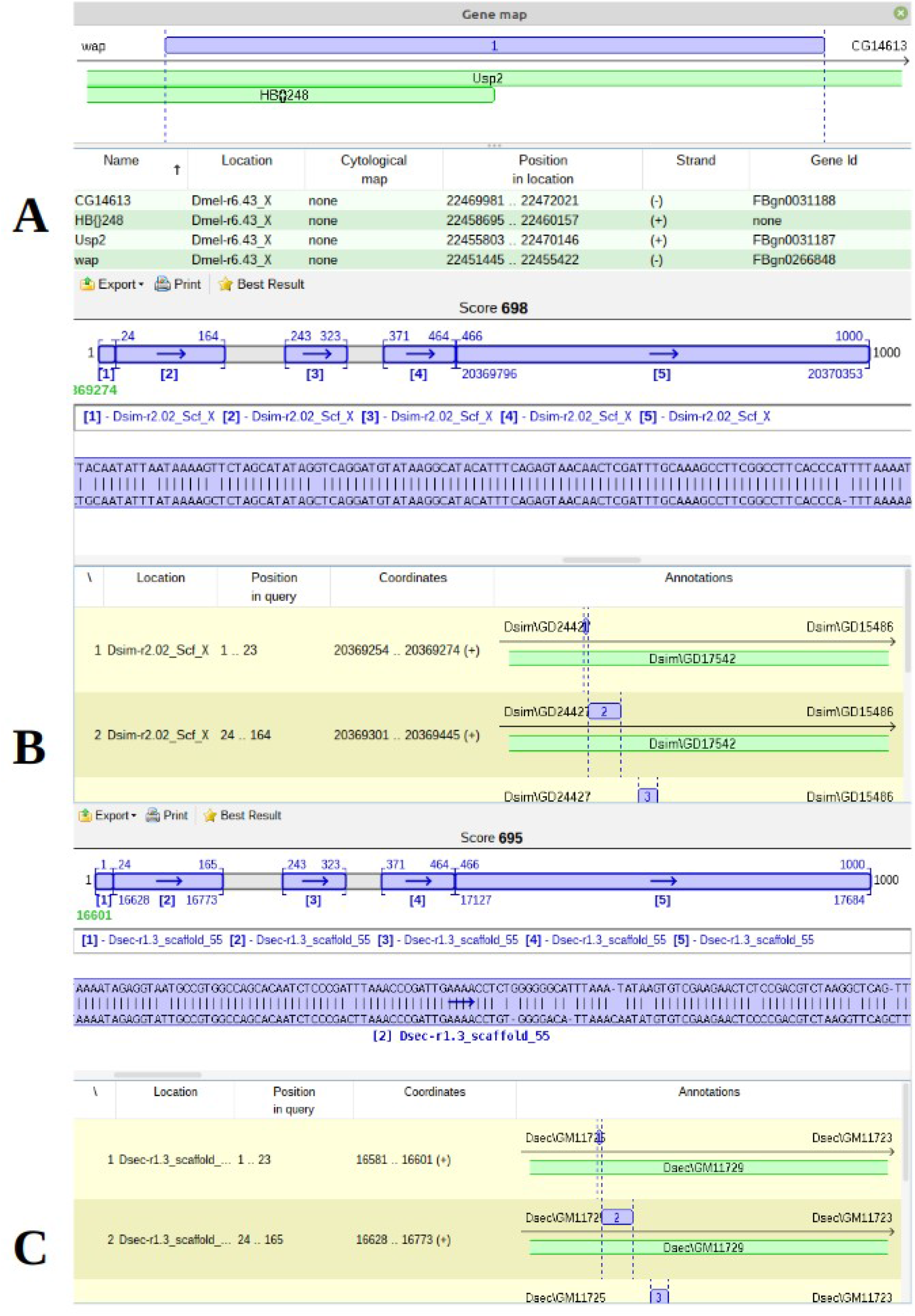
Junction sequence compose by transposon (HB{}248) and the adjacent genomic sequence.. (A) Panel A shows that this junction sequence is found in Usp2 from *D*. *melanogaster*. (B) Panel B shows that the junction sequence is identified in the GD15486 gene from *D*. *simulans*. (C) Panel C shows that the junction sequence is identified in the GM11729 gene from *D*. *sechellia*.

## Materials and Methods

For this study, we used a computer equipped with Linux Mint, an I5-1135G7 processor, 16 GB of RAM and 512 GB of SSD. The developed workflow uses the following software: *SeqKit, Genome ARTIST* version 2.0, *BLASTn* and *Kablammo*.

The comparative analysis of the sequences of interest was carried out by using *Genome ARTIST* version 2.0, developed for mapping transposons in target genomes (Ecovoiu et al., 2020). SeqKit is used to manipulate *.fasta* files. We used this toolkit to extract subsequences and to generate the reverse complement strand of a DNA sequence (Shen et al., 2016). Kablammo is a web application, used for graphical visualization of the alignments obtained with *BLASTn* (Wintersinger and Wasmuth, 2014). *BLASTn* is dedicated to identify similarities between nucleotide sequences (Altschul et al., 1990) and was run using the command line options. The genomes of the species of interest and the residual sequences of the transposons were downloaded from FlyBase (http://flybase.org/). The scripts are available in GitHub at https://github.com/DenisaConstantin/A_bioinformatics_pipeline_for_transposons_mapping.

## Conclusions

The bioinformatics pipeline described herein can assess the evolutionary kinship of some species of interest using the criterion of preserving the location of class II transposons in the respective genomes. The distribution of the location of the transposon insertions in different genomes may constitute an additional criterion used to assess or confirm the evolutionary kinship of those species.

## References

1. Altschul, S. F., Gish, W., Miller, W., Myers, E. W., & Lipman, D. J. (1990). Basic local alignment search tool. Journal of molecular biology, 215(3), 403–410. https://doi.org/10.1016/S0022-2836(05)80360-2.

2. Drosophila 12 Genomes Consortium. Evolution of genes and genomes on the *Drosophila* phylogeny. Nature 450, 203–218 (2007). https://doi.org/10.1038/nature06341.

3. Ecovoiu, Al. A., Ghita, I. C., Chifiriuc, D. I. M., Ghionoiu, I. C., Ciuca, A. M., Bologa, A. B., Ratiu, A. C. (2020). Genome ARTIST_v2 software – a support for annotation of class II natural transposons in new sequenced genomes. Cold Spring Harbor Laboratory. BioRxiv. https://doi.org/10.1101/2020.10.30.360610.

4. Gramates, L.S.; Agapite, J.; Attrill, H.; Calvi, B.R.; Crosby, M.A.; Dos Santos, G.; Goodman, J.L.; Goutte-Gattat, D.; Jenkins, V.K.; Kaufman, T.; et al. FlyBase: a guided tour of highlighted features. Genetics 2022, 220, doi:10.1093/genetics/iyac035.

5. Miller, D. E., Staber, C., Zeitlinger, J., & Hawley, R. S. (2018). Highly contiguous genome assemblies of 15 drosophila species generated using nanopore sequencing. G3 Genes|Genomes|Genetics, 8(10), 3131–3141. https://doi.org/10.1534/g3.118.200160.

6. Pevsner, J. (2015). Introduction. In: Bioinformatics and functional genomics, Bell, L., Carden, C., Dufour, B., Rowan, E., Seymour, F., A. Tan, R. Wade (eds), John Wiley & Sons, 3–16.

7. Shen, W., Le, S., Li, Y., & Hu, F. (2016). SeqKit: A cross-platform and ultrafast toolkit for FASTA/Q file manipulation. PLOS ONE, 11(10), e0163962. https://doi.org/10.1371/journal.pone.0163962.

8. Wintersinger, J. A., & Wasmuth, J. D. (2014). Kablammo: An interactive, web-based BLAST results visualizer. Bioinformatics, 31(8), 1305–1306. https://doi.org/10.1093/bioinformatics/btu808.

